# Contrasting genes conferring short and long-term biofilm adaptation in *Listeria*

**DOI:** 10.1101/2023.06.22.546149

**Authors:** William Monteith, Ben Pascoe, Evangelos Mourkas, Jack Clark, Maliha Hakim, Matthew D. Hitchings, Noel McCarthy, Koji Yahara, Hiroshi Asakura, Samuel K. Sheppard

**Author notes:** Authors to which correspondence should be addressed.

## Abstract

*Listeria monocytogenes* is an opportunistic food-borne bacterium that is capable of infecting humans with high rates of hospitalisation and mortality. Natural populations are genotypically and phenotypically variable, with some lineages being responsible for most human infections. The success of *L. monocytogenes* is linked to its capacity to persist on food and in the environment. Biofilms are an important feature that allow these bacteria to persist and infect humans, therefore, understanding the genetic basis of biofilm formation is key to understanding transmission. We sought to investigate the biofilm forming ability of *L. monocytogenes* by identifying genetic variation that underlies biofilm formation in natural populations using genome-wide association studies. Changes in gene expression of specific strains during biofilm formation were then investigated using RNAseq. Genetic variation associated with enhanced biofilm formation was identified in 273 genes by GWAS and differential expression in 220 genes by RNAseq. Statistical analyses show that number of overlapping genes flagged by either type of experiment is less than expected by random sampling. This is consistent with an evolutionary scenario where rapid adaptation is driven by variation in gene expression of pioneer genes, and this is followed by slower adaptation driven by nucleotide changes within the core genome.

**Impact statement:** *Listeria monocytogenes* is a problematic food-borne bacterium that can cause severe illness and even death in humans. Some strains are known to be more common in disease and biofilms are crucial for survival in the environment and transmission to humans. To unravel the genetic basis of biofilm formation, we undertook a study employing genome-wide association studies (GWAS) and gene transcription profiling. We identified 273 genes associated with robust biofilm formation through GWAS and discovered differential expression in 220 genes through RNAseq. Statistical analysis revealed fewer overlapping genes than expected by chance, supporting an evolutionary scenario where initial adaptation relies on gene expression variation, followed by slower adaptation through genetic changes within the core genome.

**Data summary:** Short read genome data are available from the NCBI (National Center for Biotechnology Information) SRA (Sequence Read Archive), associated with BioProject PRJNA971143 (https://www.ncbi.nlm.nih.gov/bioproject/PRJNA971143). Assembled genomes and supplementary material are available from FigShare: doi: 10.6084/m9.figshare.23148029. RNA sequence data and differential gene expression profiles have been deposited in the NCBI Gene Expression Omnibus.

## Introduction

*Listeria monocytogenes* is an opportunistic non-fastidious bacterium that causes serious invasive disease in humans and animals, known as listeriosis (Farber and Peterkin, 1991; Murray, Webb and Swann, 1926). Infection with *L. monocytogenes* is caused primarily by the consumption of contaminated food products, with infections displaying the highest hospitalisation rate of any food-borne pathogen, at 92% of cases (Mead *et al*., 1999), and a high mortality rate, at 20% to 30% of cases (Pizarro-Cerdá and Cossart, 2019). In fact, *L. monocytogenes* is responsible for the third most deaths caused by any food-borne pathogen in the United States, despite infections only accounting for a small minority of food-borne disease cases. This problem is only expected to increase, as cases of *L. monocytogenes* have continued to rise in North America and Europe over the last decade (EFSA and ECDC, 2018; Scallan *et al*., 2011).

*L. monocytogenes* is commonly isolated from animal hosts (cattle, sheep and birds) (Mpundu *et al*., 2022; Rothrock *et al*., 2017; Hellström *et al*., 2008; Grønstøl, 1980) and their food products (raw meat, pasteurised milk, fish) (Kayode and Okoh, 2022; Osman *et al*., 2020; Osman *et al*., 2014), as well as environmental sources, such as soil, vegetables, sewage and other water sources (Mpondo, Ebomah and Okoh, 2021; Zhu, Gooneratne and Hussain, 2017; Vivant *et al*., 2015; Linke *et al*., 2014; Ramaswamy *et al*., 2007). Consumption of any of these food sources can lead to infection of the host via translocation of the intestinal barrier, infection of the lymph nodes, and the pathogen subsequently entering the spleen and liver (Werbrouck *et al*., 2006). Infection of healthy individuals frequently results in febrile gastroenteritis, a non-invasive disease manifesting within a day of ingesting contaminated food (Dalton *et al*., 1997). However, in immunocompromised patients unrestricted proliferation of *L. monocytogenes* can cause bacteremia, leading to invasion of additional organs, especially the brain and spinal cord, causing meningitis (Longhi *et al*., 2004).

The population structure of *L. monocytogenes* reveals four phylogenetic lineages that are distinct in terms of their pathogenicity and host specificity. Lineages I and II are associated with food and human outbreak samples whereas lineages III and IV are found in a broader range of environments and are usually isolated from animal sources (Orsi, den Bakker and Wiedmann, 2011). Isolates of lineage I have been most isolated from epidemic listeriosis samples whereas lineage II is associated with food and related food production environments (Moura *et al*., 2021; Nightingale *et al*., 2008). There are several hundred clonal complexes (CCs) identified for *L. monocytogenes*. Historically CC1 and CC2 have been the most prevalent isolates (Ragon *et al*., 2008). However, in recent years CC1 and CC2 are found less often in datasets and previously non-typical isolates, especially CC5, CC6, and CC121, are rising in frequency (Bergholz *et al*., 2018). The Listeria serotyping scheme has also identified 13 serotypes of *L. monocytogenes* (Ragon *et al*., 2008; Palumbo *et al*., 2003). Most human cases of listeriosis are sporadic and tend to involve isolates of serotypes 1/2a, 1/2b and 4b, with serotype 4b being an important contributor (∼36% of cases) (Lee *et al*., 2014; Swaminathan and Gerner-Smidt, 2007; Gasanov, Hughes and Hansbro, 2005).

Multicellular biofilm formation is a key feature for bacterial tolerance to adverse environmental conditions (López, Vlamakis and Kolter, 2010)(Otto, 2014). In *L. monocytogenes,* biofilms promote surface adherence (Lee, Hébraud and Bernardi, 2017; Møretrø and Langsrud, 2004) and persistence in food production environments (Rodríguez-Campos *et al*., 2019; Ulusoy and Chirkena, 2019; Sofos and Geornaras, 2010; Greenwood, Roberts and Burden, 1991), even enabling survival and replication in refrigeration conditions (Bucur *et al*., 2018)(Chan and Wiedmann, 2009; Becker *et al*., 2000) and resistance to antimicrobials, including disinfectants (Wiśniewski *et al*., 2022; Wiktorczyk-Kapischke *et al*., 2021; Skowron *et al*., 2019; Conficoni *et al*., 2016). This is a major public health concern since food contamination can result in fatal listeriosis outbreaks following consumption (McLauchlin, Grant and Amar, 2020; Meldrum *et al*., 2009). Interestingly, biofilm forming strains are structurally and physiologically different from their planktonic counterparts, with the former being more coccoid than rod shaped, and displaying increased antibiotic resistance. Furthermore, biofilms formed by *L. monocytogenes* strains found in food industries are thicker than those formed by sporadically occurring isolates (Colagiorgi *et al*., 2016). Therefore, there is clear incentive to understand the genetics of biofilm formation.

Genes involved in cell motility, surface adhesion, and expression of extracellular polymers are all linked to *L. monocytogenes* biofilm formation (Popowska *et al*., 2017; Colagiorgi *et al*., 2016; Lemon, Higgins and Kolter, 2007). However, biofilms are a complex phenotype that can be underpinned by numerous micro-evolutionary changes to the genome operating over different timescales. For example, short-term genetic changes may include mutations affecting gene expression such as phase variation, rapid and reversibly to repeat regions can switch a particular gene between expression states (Aidley *et al.,* 2018)(Brooks and Jefferson, 2014; Levinson and Gutman, 1987). Phase variation within the *inlA* gene has been implicated as a mechanism of niche adaptation and virulence attenuation in *L. monocytogenes* (Manuel *et al*., 2015b). In contrast to rapid adaptation and expression state switching, *de novo* mutation and horizontal gene transfer (HGT), usually considered infrequent in *L. monocytogenes,* can also introduce genetic variation conferring novel functionality (Ragon *et al*., 2008)(den Bakker *et al*., 2008). However, these processes take place over long timescales than phase variation.

High-throughput sequencing and growing collections of sequenced isolates, provide opportunities to investigate the genetic basis of complex traits like biofilm formation. Comparative transcriptomics and genome-wide association studies (GWAS) provide an effective method to identify associations between genetic variants and phenotypes (Uffelmann *et al*., 2021; Sheppard *et al*., 2013). In this study we use a quantitative population genomics approach to investigate the genes involved in short and long-term adaptation to biofilm formation in *L. monocytogenes*.

## Methods

### Bacterial sampling

The isolate collection comprised 868 *Listeria monocytogenes* isolates from a variety of sources (**Table S1)**. These included 760 publicly available whole genome sequences from the Institut Pasteur database (Moura *et al*., 2016; Jolley and Maiden, 2010) and 108 isolates sampled in Japan and sequenced as part of this study. Of the Japanese samples, 41 were sampled from food products, specifically 10 from chicken, 9 from beef, 18 from pork, 3 from salad products and 1 from fish, 35 isolates were sampled from agricultural sources (cow and colostrum), 28 isolates were sampled from human clinical cases, and 4 isolates were from environmental samples.

### Genomic DNA extraction and whole genome sequencing

*L. monocytogenes* isolates were cultured in Todd-Hewitt broth at 37°C for 18-24 hours, and genomic DNA was extracted using the QIAmp DNA Mini Kit (QIAGEN, Crawley, UK), according to manufacturer’s instructions. DNA was quantified using a Nanodrop spectrophotometer before sequencing. Genome sequencing was performed on an Illumina MiSeq sequencer (Illumina, Cambridge, UK) using the Nextera XT Library Preparation Kit with standard protocols. Libraries were sequenced using 2 x 300 bp paired end v3 reagent kit (Illumina, following manufacturer’s protocols). Short read paired-end data were filtered, trimmed, and assembled using the *de novo* assembly algorithm, SPAdes (version 3.7) (Prjibelski *et al*., 2020). The average number of contigs in 108 *L. monocytogenes* genomes was 564 (range: 37-3790) for an average total assembled sequence size of 2.91 Mbp (range: 2.00-3.71). The average N50 was 109313 (range: 958-583650) and the average GC content was 38.2% (range: 37.3-43.0). An overview of assembly information is provided in **Table S1**. Genomes and short read data are archived on the NCBI GenBank and SRA depositories, associated with BioProject accession #PRJNA971143.

### Phenotype testing

*L. monocytogenes* is commonly isolated from diverse environmental sources and can persist in harsh conditions. To understand this persistence, we sought to quantify aspects of biofilm formation. Biofilm formation was measured by a semi-quantitative adherence assay using 96– well tissue culture plates (Pascoe *et al*., 2015; Mack *et al*., 1994). 5 μl aliquots of overnight culture were used to inoculate 200 μl of liquid media and plates were then incubated on a moving platform at either 30°C or 37°C for 48 h. Culture medium was removed and wells washed with PBS. Bacteria adhered to the plate were fixed with 150 μl of Bouin’s solution before being washed with PBS. Plates were left to dry and stained with 150 μl of 0.1% (w/v) crystal violet for 5 min. Spectrophotometric measurements were taken at OD600 in a 96– well plate reader and the average of three replicates was calculated (BMG Omega).

### Pangenome archiving and phylogenetic tree construction

Whole genome assemblies were annotated using Prokka (version 1.12) (Seemann, 2014). Annotated assemblies were processed by the PIRATE pan-genome pipeline (version 1.0.4) (Bayliss *et al*., 2019) to identify clusters of orthologous genes (COGs). Gene sequences were translated to amino acid sequences and COGs were defined by PIRATE using a wide range of amino acid percentage sequence identity thresholds for Markov Cluster algorithm (MCL) clustering (45, 50, 60, 70, 80, 90, 95, 98). The pangenome of all 868 *L. monocytogenes* isolates contained 23637 COGs. The pangenome of 108 *L. monocytogenes* isolated in Japan contained 11202 COGs. Genes shared by >95% of the isolates were defined as part of the core genome. Concatenated core genome alignments were produced gene-by-gene using MAFFT (All isolates (*n*=868): 1642 genes, 1656306 bp; Japanese isolates (*n*=108): 869 genes, 844305 bp) and the phylogenies were inferred from these alignments using a maximum likelihood (ML) algorithm using RAxML (version 8.2.4) (Stamatakis, 2014) with the generalized time-reversible (GTRGAMMA) substitution model. The resulting phylogenies and core-genome alignments were used as input to ClonalFrameML (version 1.12) (Didelot and Wilson, 2015) to account for recombination taking place within the core-genome.

### RNA preparation for RNA-seq

To understand changes in gene expression involved in biofilm formation, we performed a differential gene expression analysis of *L. monocytogenes* in a planktonic state, whereby bacteria were grown in shaken media to prevent the formation of any biofilms, versus bacteria in a biofilm forming state. The transcriptomes of two isolates, Lm0132 and Lm0134, were analyzed. These belong to two of the most common *L. monocytogenes* complexes, CC2 and CC1, respectively. Strains were cultured in Todd-Hewitt broth at 37°C for 24 hours. RNA was isolated using the SV RNA isolation kit (Promega) according to the manufacturer’s instructions. 23S and 16S rRNA was depleted using a MicrobExpress kit (Ambion) and genomic DNA digested using amplification-grade DNase I (Invitrogen). RNA was reverse-transcribed using random primers (Invitrogen) and Superscript III (Invitrogen) at 45 °C for 3 h and denatured at 70 °C for 15 min. Nextera XT libraries were constructed and sequenced with over 30,000 reads per sample (min estimated coverage above x107).

### Differential gene expression analysis

Illumina sequence reads containing expression data for each of the two strains, and two independent RNA samples for each condition, were mapped to a reference transcriptome index created using Salmon (version 1.9.0) against a genome reference F2365 (Patro *et al*., 2017). Gene expression was calculated in units of transcripts per million (TPM), a relative abundance measure used for downstream analysis. DESeq2 in Bioconductor (Huber *et al*., 2015; Love, Huber and Anders, 2014) was used for differential expression analysis. Briefly, raw counts were normalized using size factors to account for differences in library depth. Counts were further log-transformed using the variance stabilizing transformation function. Gene-wise dispersion was calculated, and the estimates were reduced using empirical Bayesian shrinkage to avoid biological variance. Hypothesis testing was applied using the Wald’s test across the negative binomial distribution. Differences in fold-change values were calculated by determining the log2 fold-change (LFC). A stringent cut-off of LFC <-2 and/or >2 was used to identify differentially expressed genes. An adjusted *p*-value was calculated for every gene with those *p*<0.05 being considered significant. Initial testing in DESeq2 found that expression data for isolate Lm0134 was of poor quality. Reads from Lm0132 strain taken from the 24 h shaken media were compared with reads taken from 24 h stationary growth.

### Genome-wide association studies

GWAS was performed to test for genetic variation associated with biofilm formation. Genetic variation within the *L. monocytogenes* pangenome was covered by extracting variable length *k*-mers from the assemblies, known as unitigs. Pyseer (version 1.3.6) (Lees *et al*., 2018) was used to fit an elastic net model to test the association between unitigs and biofilm formation at both 30°C and 37°C. The elastic net uses regularized linear regression to simultaneously test the association of all the genetic variants with the phenotype and reports the variants predicted to be causal for the phenotype (Lees *et al*., 2020). The lasso default correlation filter of 25% was used. Only unitigs with a minor-allele frequency (MAF) of >2% were selected. 872250 unitig sequences were generated from the *L. monocytogenes* pangenome. 436125 filtered variants were removed from the analysis. 436125 variants were tested for their association with the phenotype. Despite the elastic net’s implicit control for the effects of population structure by distinguishing genetic variants that are causal for the phenotype from those inherited from the common ancestor (lineage effects), an explicit correction was also applied by using a genetic relatedness matrix calculated from the inferred population phylogeny to adjust the p-value of association variants. The phylogenetically corrected p-values were used by a suite of tools to interpret the contribution of genetic variation highlighted by the model. To infer gene function, unitigs were mapped to the *L. monocytogenes* F2365 complete reference genome (assembly accession: NC_002973.6) using BWA-MEM (version 0.7.17) (Li, 2013).

### Gene function determination

To determine gene function the KEGG database was used (Kanehisa *et al*., 2023; Kanehisa *et al*., 2016). Briefly, custom python scripts were employed to interface with the database by iteratively querying all *L. monocytogenes* schemes until a match was identified. KEGG information was downloaded via the KEGG database API and the resulting text file was parsed and gene functions extracted. Where a gene had more than one function, all functions were extracted.

## Results

### L. monocytogenes populations are highly structured

We investigated the epidemiology and whole-genome sequences of *L. monocytogenes* isolated from multiple sources in Japan. A total of 108 isolates were retrieved from diverse sources, including clinical, food production and agricultural sources. These isolates were tested for their biofilm production and underwent whole-genome sequencing (**Table S2**). In addition, 760 isolates from public databases were included in our analyses to provide context and an indication of the natural population structure of *L. monocytogenes* (**Table S1**)

Core-genome phylogenetic analysis revealed the population structure of *L. monocytogenes* (**Figure 1**). The maximum-likelihood core-genome phylogeny was highly structured and included the major phylogenetic lineages I, II, III and IV. In total the dataset consisted of 277 unique sequence types (STs), of which 152 belong to 82 clonal complexes and 125 of which did not belong to a previously defined complex. Despite our collection of samples all being sourced from Japan, the 108 isolates are representative of the diversity shown in the clinically relevant lineages I and II and were classified into 33 STs from 27 clonal complexes. However, the addition of 760 publicly available samples revealed increased diversity, due the addition of 137 lineage III isolates, which are not observed in the Japanese isolate collection. Lineage III is the most genetically diverse lineage and included 124 STs, of which 21 belonged to 16 clonal complexes, and 103 of which did not belong to a previously defined clonal complex. Furthermore, the additional lineage I and II isolates also included STs not present in our Japanese collection, with 38 and 80 new STs being added for each, respectively.

**Figure 1:**
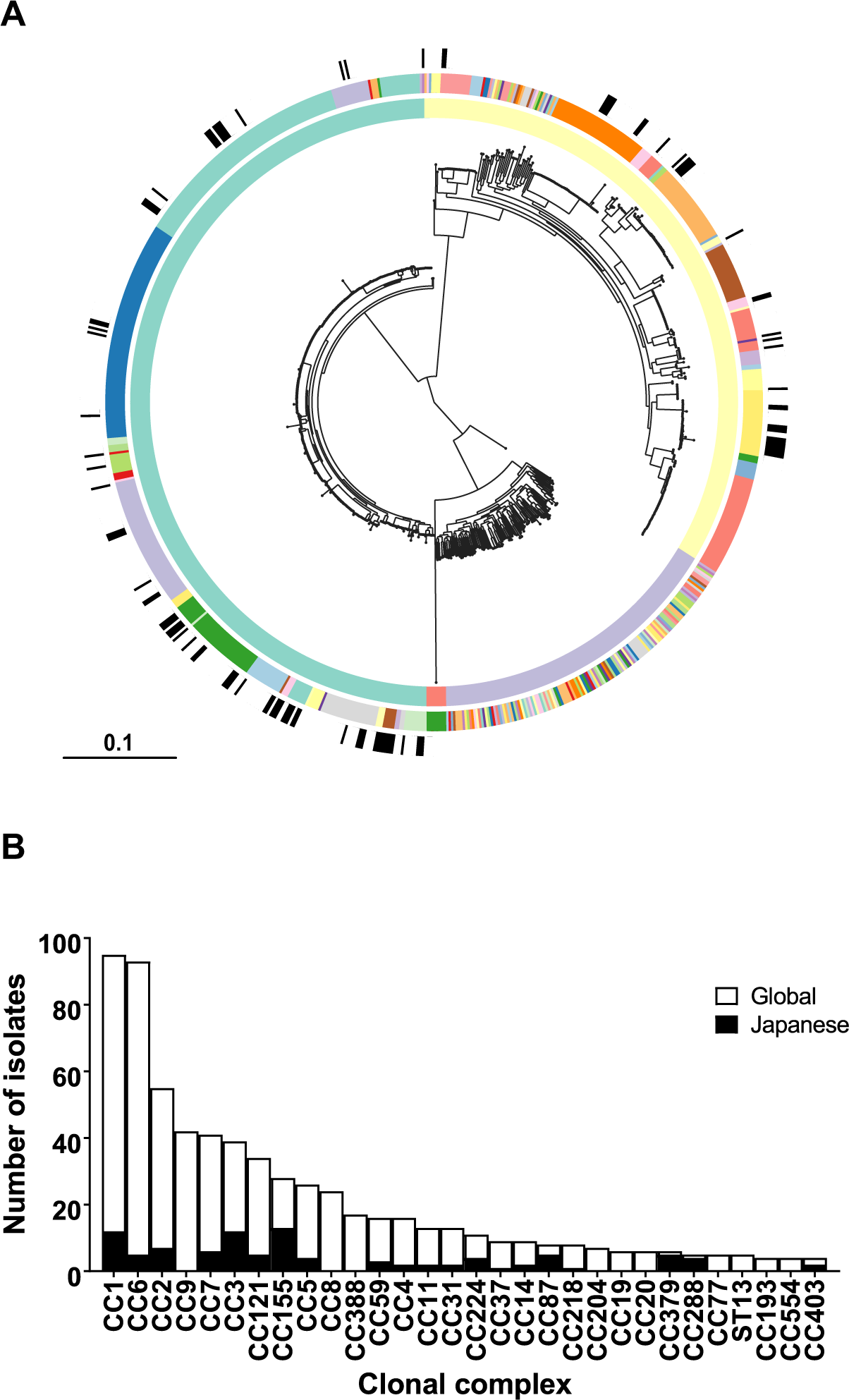
*L. monocytogenes* clones isolated in Japan are globally distributed. (A) *L. monocytogenes* isolates (*n*=868) are shown on a core-genome phylogenetic tree reconstructed using a ML algorithm implemented in RAxML. The rings, from inner to outer, indicate the phylogenetic lineage of isolates (lineage I is green, lineage II is yellow, lineage III is purple, lineage IV is red); the diversity of clonal complexes present within each lineage; the distribution of Japanese isolates across the phylogeny is indicated in black. (B) The frequency of *L. monocytogenes* isolates by clonal complex. Only complexes with four or more isolates are included. Bars are coloured according to isolation location (Japan is black, the rest of the world is white).

### Biofilm formation varies among L. monocytogenes isolates

The extent of biofilm formation was measured at 30°C and 37°C from the absorbance value (OD600) of adhered cells following fixation and staining with crystal violet (**Figure 2**). Absorbance values recorded at 30°C were significantly greater than at 37°C with a mean reading for biofilm thickness of 0.489 (range: 0.242-0.742) and 0.413 (range: 0.208-0.671), respectively (P<0.0001). Of the 108 isolates studied, 78.7% (*n* = 85) produced more biofilm at 30°C than at 37°C.

**Figure 2:**
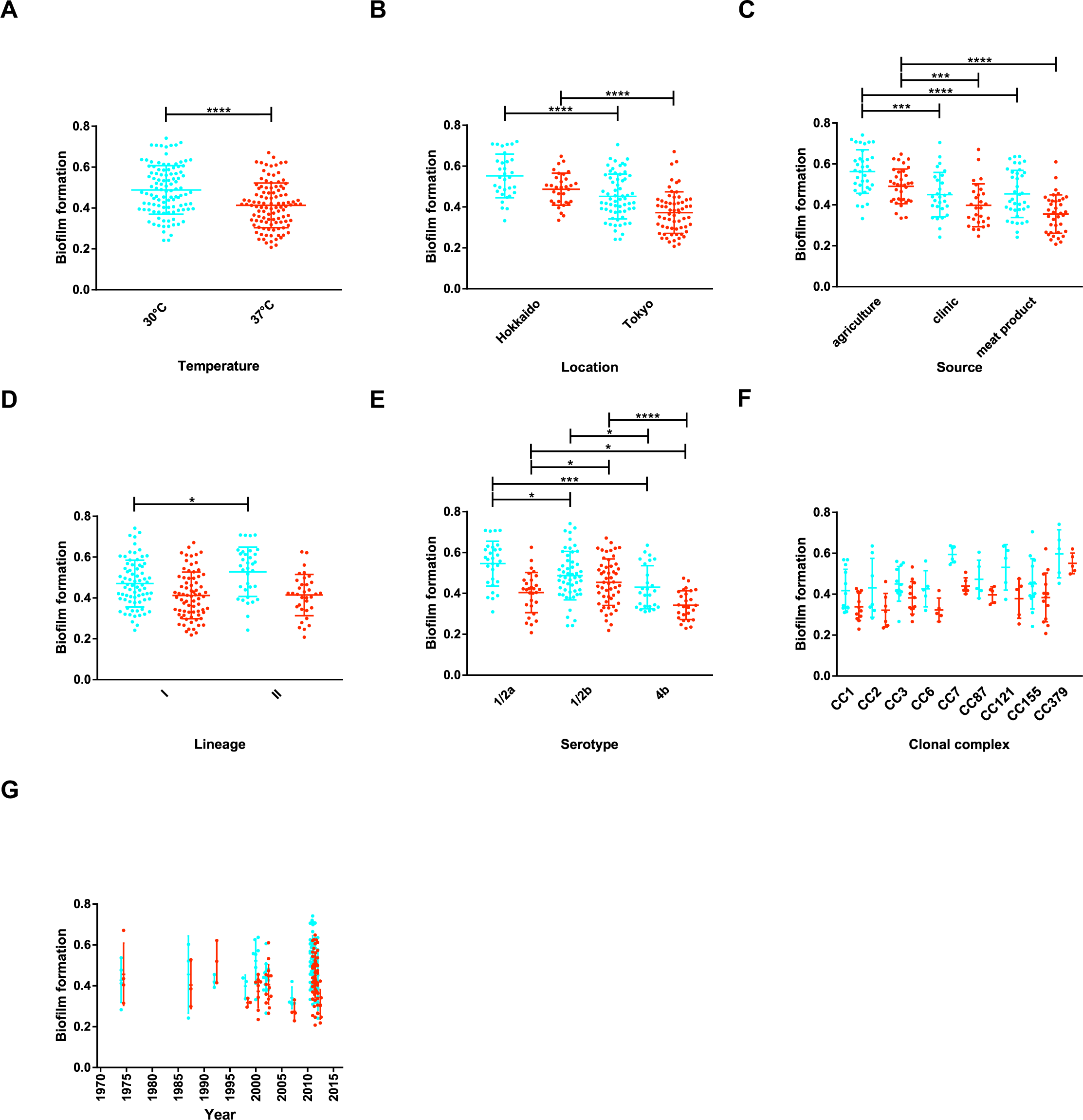
Epidemiological factors associated with biofilm formation in Japan. Box plots showing the association between biofilm forming ability and (A) temperature, (B) isolation location, (C) isolation source, (D) phylogenetic lineage, (E) serotype, (F) clonal complex, and (G) time. Statistical analysis was performed by Student’s *t* test comparing biofilm thickness in each condition (\**p* < 0.05; \*\**p <* 0.01; \*\*\**p* < 0.001; \*\*\*\**p* < 0.0001).

Enhanced biofilm forming ability was associated with multiple isolation sources. Isolates sampled from agricultural sources had significantly greater biofilm thickness than isolates sampled from clinical sources and meat products at both temperatures. Biofilm formation was also significantly greater in isolates sampled in Hokkaido, which consisted of samples from agricultural sources (*n*=28) and very few clinical samples (*n*=2) compared with those isolated in Tokyo, which mostly consisted of isolates sampled from the clinic (*n*=22) and meat production environments (*n*=35) and did not include any agricultural samples.

Enhanced biofilm forming ability was observed in isolates of lineage II compared with isolates of lineage I at 30°C, however, this difference in biofilm thickness was not observed between lineages at 37°C. Further investigation, of the biofilm forming ability by clonal complex revealed an unequal distribution; isolates belonging to CC7 and CC379 had significantly greater biofilm thickness than other complexes. The fact that CC7 belongs to lineage II, whilst CC379 belongs to lineage I might explain why the difference in biofilm forming ability between the phylogenetic lineages was not consistent between temperatures.

Biofilm formation was also compared among isolates from the most prevalent serotypes (1/2a, 1/2b and 4b). Serotypes 1/2a and 1/2b showed significantly greater biofilm thickness compared with 4b isolates at both temperatures. However, it was less clear whether there was a difference in biofilm forming ability between serotypes 1/2a and 1/2b, with 1/2a isolates showing greater biofilm thickness at 30°C and 1/2b showing greater biofilm thickness at 37°C.

### L. monocytogenes biofilm formation is not explained by phylogenetic structure

A maximum-likelihood phylogeny was constructed from the core-genome alignment of 108 Japanese isolates and the biofilm forming ability at 30°C and 37°C was mapped onto the tree (**Figure 3**). The phylogeny revealed that biofilm forming ability varied across the *L. monocytogenes* population, with high biofilm forming strains occupying divergent phylogenetic positions. Except for CC7 and CC379, biofilm forming ability was not strain associated. Furthermore, there was a positive correlation between the biofilm forming ability of isolates at 30°C and 37°C, indicating that biofilm formation was independent of temperature across this range, i.e., individual isolates capable of forming thick biofilms at 30°C also formed thick biofilm at 37°C. This evidence implies that there are common genomic determinants underlying *L. monocytogenes* biofilm formation.

**Figure 3:**
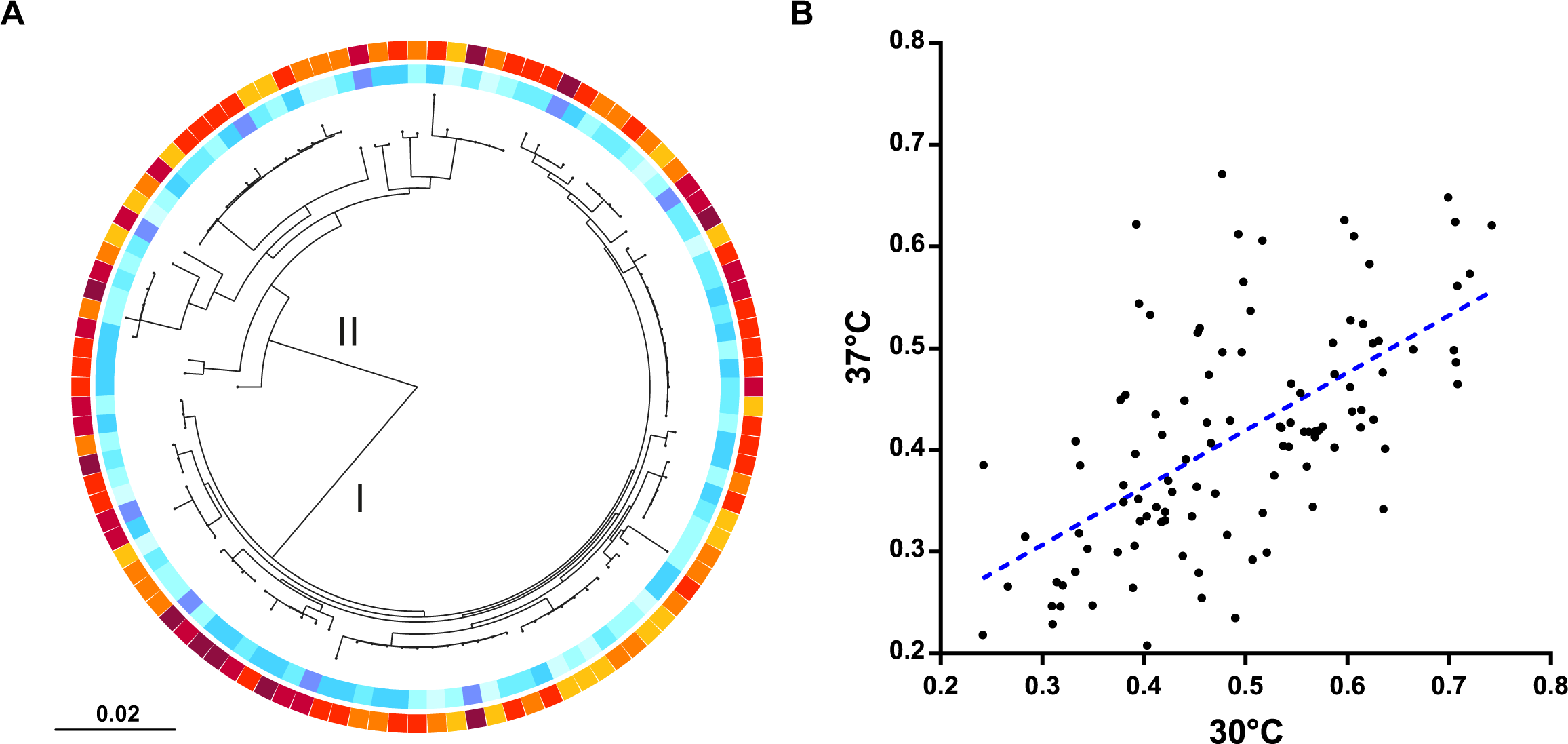
Biofilm formation is distributed throughout the clonal frame of *L. monocytogenes*. (A) *L. monocytogenes* isolates (*n*=108) from Japan are shown on a core-genome phylogenetic tree reconstructed using a ML algorithm implemented in RAxML. The coloured rings indicate the biofilm forming ability of isolates at 30°C (inner) and 37°C (outer). The scale bar indicates the estimated number of substitutions per site. (B) Scatter plot of isolates biofilm formation at 30°C and 37°C. The blue dotted line indicates the line of best fit.

### Genome-wide association study identifies genetic elements associated with biofilm formation

The genetic association of enhanced biofilm formation in *L. monocytogenes* was investigated at 30°C and 37°C in 108 Japanese isolates demonstrating a range of biofilm forming phenotypes (**Figure 4**). A maximum-likelihood phylogeny of the 108 isolates was inferred using a core-genome alignment and 436125 variable length unitig *k*-mers generated from the pangenome of these isolates were tested for their association with the biofilm formation phenotype. 4079 and 4019 unitigs were identified by GWAS to be associated with biofilm formation at 30°C and 37°C, respectively. A Bonferroni corrected significance threshold of – log_10_ (*P*) = 6.97 was calculated as the genome-wide significance level, and the suggestive significance threshold of –log_10_ (*P*) = 3.97 was used. To reveal genetic regions and genes associated with biofilm formation, *k*-mers were mapped to the reference *L. monocytogenes* F2365 genome (**Table S3 & S4**). The KEGG database resource API (Kanehisa *et al*., 2023; Kanehisa *et al*., 2016) was used to derive gene functions from the *L. monocytogenes* database (**Table S5**).

**Figure 4:**
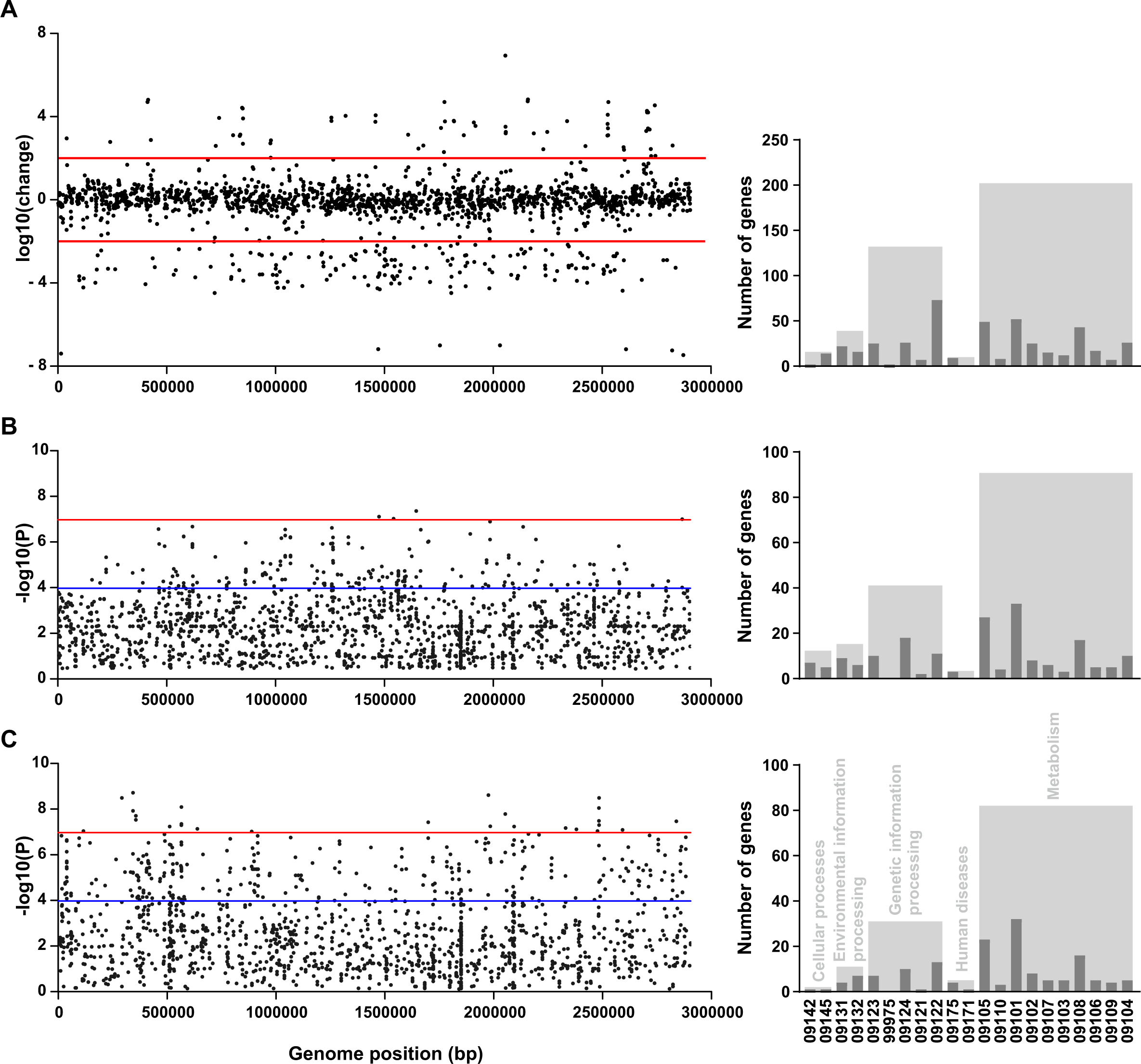
Genes with differential expression in transcription analysis are distinct to genes associated with biofilm formation in genome-wide association studies. (A) Differential gene expression between planktonic and biofilm forming state of Lm0132. The LOG-fold change in gene expression against its position in the F2365 genome is shown. Red lines denote a significance threshold of 100x change in gene expression. (B) & (C) Manhattan plots of unitigs associated with biofilm formation at 30°C and 37°C, respectively. The significance of each unitig *k*-mer’s association with biofilm formation against its position in the *L. monocytogenes* F2365 genome is shown. The red line denotes genome-wide significance threshold –log_10_ (*P*) = 6.97 (α < 0.05 Bonferroni corrected with 436125 unique variants), and the blue line denotes the suggestive significance threshold –log_10_ (*P*) = 3.97. Bar charts represent the number of genes exceeding significance thresholds with a function described in the KEGG database (Kanehisa *et al*., 2016). Dark grey bars represent gene functions (09142: Cell motility, 09145: Cellular community – prokaryotes, 09131: Membrane transport, 09132: Signal transduction, 09123: Folding, sorting and degradation, 99975: Protein processing, 09124: Replication and repair, 09121: Transcription, 09122: Translation, 09175: Drug resistance – antimicrobial, 09171: Infectious disease – bacterial, 09105: Amino acid metabolism, 09110: Biosynthesis of other secondary metabolites, 09101: Carbohydrate metabolism, 09102: Energy metabolism, 09107: Glycan biosynthesis and metabolism, 09103: Lipid metabolism, 09108: Metabolism of cofactors and vitamins, 09106: Metabolism of other amino acids, 09109: Metabolism of terpenoids and polyketides, 09104: Nucleotide metabolism), light grey bars represent functional classes (09140: Cellular Processes, 09130: Environmental Information Processing, 09120: Genetic Information Processing, 09160: Human Diseases, 09100: Metabolism).

At 30°C there were 5 biofilm-associated *k*-mers exceeding the genome-wide significance threshold and 230 biofilm-associated *k*-mers exceeding the suggestive significance threshold. These *k*-mers mapped to 3 and 114 genes, respectively (**Table S3**). At 37°C there were 46 biofilm-associated *k*-mers exceeding the genome-wide significance threshold and 462 biofilm-associated *k*-mers exceeding the suggestive significance threshold. These *k*-mers mapped to 18 and 181 genes, respectively (**Table S4**). In total, 676 unique *k*-mers were found to be significantly associated with biofilm formation, these mapped to 273 genes, including 22 genes shared by both 30°C and 37°C conditions. These included genes that have previously been shown to have a functional association with biofilm formation, such as cell wall or cell membrane biogenesis (Bucher *et al*., 2015), motility (Lemon, Higgins and Kolter, 2007) and protein glycosylation (Iwashkiw *et al*., 2012). Additionally, genes with the potential to influence biofilm formation in *Listeria monocytogenes* included genes putatively involved in energy production and conversion, transport and metabolism of amino acids, carbohydrates, lipids and nucleotides (Greenwich *et al*., 2019; Kolodkin-Gal *et al*., 2010; Lu *et al*., 2019; Harmsen *et al*., 2010).

Genes identified by GWAS have diverse roles. Multiple genes involved in the biosynthesis of amino acids were found to be associated with biofilm formation at both temperatures, suggesting that biofilms may increase the production of enzymes involved in producing scarce amino acids. These include *aroC* and *aroE*, which are involved in the synthesis of aromatic amino acids via the shikimate pathway. AroC has previously been demonstrated to affect biofilm formation in *Escherichia coli*, *Streptococcus mutans* and *Edwardsiella tarda*, and is essential for the production of intermediates, such as indole, that affect flagellar synthesis and biofilm formation (Liu *et al*., 2022; Puttamreddy, Cornick and Minion, 2010; Shemesh, Tam and Steinberg, 2007). AroE has also previously been found to be associated with biofilm adaptation in *L. monocytogenes* (Santos *et al*., 2019). Another important mechanism by which genes involved in amino acid metabolism may be utilized in biofilm adaptation includes stress tolerance. The transcriptional repressor *argR* is involved in arginine biosynthesis and is induced by the central virulence regulator PrfA, known to have an important role in *L. monocytogenes* capacity to withstand low pH (Ryan *et al*., 2009). Furthermore, argR has previously been demonstrated to affect the biofilm forming ability of *Streptococcus gordonii* (Robinson *et al*., 2018; Jakubovics *et al*., 2015). Other genes involved in amino acid metabolism that were flagged include *gcvT* (Melian *et al*., 2021), *hisJ* (Zhang *et al*., 2020), and *thrB* (Ge *et al*., 2008), which have all previously been demonstrated to affect biofilm formation in other bacterial species.

In *L. monocytogenes*, flagellum-mediated motility is critical for attachment to surfaces and subsequent biofilm formation (Lemon, Higgins and Kolter, 2007). Our findings support this as variation within genes involved in cell motility appeared to play an important role in biofilm formation. The major virulence determinant gene, *actA*, was found to be associated with biofilm formation at both temperatures. ActA has previously been shown to be critical in *L. monocytogenes* biofilm formation (Travier *et al*., 2013). In addition, the flagellar motor switch gene (*fliM*) was also identified (Melian *et al*., 2021). Furthermore, additional genes involved in cell motility were found to be associated with biofilm formation at 30°C (*fliF*, *fliR, flgE*, *flgK*, *motA* and *motB*), further emphasizing the importance of motility in the ability of *L. monocytogenes* to form biofilms.

Genes involved in cell wall or cell membrane biogenesis are essential for biofilm formation. Indeed, multiple genes flagged by the genome-wide association studies are already known to be involved in biofilm formation. The *dltB* gene has been shown by mutagenesis studies of *L. monocytogenes* to have an important role in biofilm formation (Alonso *et al*., 2014). D-alanylation of extracellular lipoteichoic acids appears to be important in maintaining an appropriate surface charge for bacterial attachment to abiotic surfaces. Similarly, *inlH* is a stress induced virulence gene which produces a surface protein that facilitates *L. monocytogenes* survival in tissues. Inactivation of *inlH* has been shown to enhance biofilm formation in *L. monocytogenes* (Janež *et al*., 2021).

### Differential expression of biofilm associated genes

The differential expression of genes involved in biofilm formation was investigated by comparing the transcriptome of *L. monocytogenes* in shaken culture versus static growth. This analysis found a total of 220 genes were differentially expressed between conditions with a log10-fold change greater than 2 and a significance value of *P*<0.05. A reduction in gene expression was seen between the planktonic state and biofilm formation phase of bacterial growth in 160 genes, whilst 60 genes increased in expression (**Table S6**). The function was determined for all genes (**Table S5**).

Genes involved in cell wall or cell membrane biosynthesis, (*dltB*, *dltC* and *galE*), had changes in gene expression associated with biofilm formation. The *dlt* genes were down-regulated in the biofilm state, whereas transcription of *galE* was up-regulated. Genes from the dltABCD operon have previously been implicated in biofilm formation in *L. monocytogenes* and other bacterial species (Alonso *et al*., 2014), and mutation within genes involved in glycosylation of the LPS has previously been shown to affect biofilm formation. A *galE* mutant of *Porphyromonas gingivalis* was shown to have enhanced auto-aggregation and biofilm formation by increasing surface hydrophobicity, however, this phenotype was only partially replicated in a *galE* mutant of *Escherichia coli* (Nakao *et al*., 2012; Nakao, Senpuku and Watanabe, 2006). Similarly, multiple genes involved in cell division showed changes in gene expression. *ftsX* and *mreB* both showed a large increase in transcription in the biofilm phase. In *Fusobacterium nucleatum,* deletion of *ftsX* abolished biofilm formation (Wu *et al*., 2018). Mutation to the *ftsEX* operon in *Bacillus veleznsis* also lead to defective biofilm formation (Li *et al*., 2018). Furthermore, the *ftsX* gene is down-regulated in *L. monocytogenes* in response to high hydrostatic pressure, suggesting the cell-division protein has a role in adaptation to stressful conditions (Osek, Lachtara and Wieczorek, 2022). The bacterial actin homolog *mreB*, which has a putative function in cell wall organization and determining cell shape, was also up-regulated in the biofilm phase (Hutchins *et al*., 2021; van Teeffelen and Gitai, 2011). c-di-AMP is a secondary messenger that fulfils multiple important functions in bacteria (Stülke and Krüger, 2020). In *L. monocytogenes* disruption of c-di-AMP metabolism due to mutant strains lacking the phosphodiesterase *PgpH* affects biofilm formation and integrity of the cell envelope (Wang *et al*., 2022; Whiteley *et al*., 2017; Peng *et al*., 2016).

Genes involved in amino acid transport and metabolism were also differentially expressed. In addition to *argH,* which was flagged by GWAS, multiple genes involved in amino acid metabolism were identified, including *aroH, asnB, dapF, hisC, purF,* and *serC*. Multiple genes involved in nucleotide metabolism or DNA replication and repair were found to be differentially expressed in *L. monocytogenes*. For example, both *ruvA* and *ruvB* displayed a reduction in expression in the biofilm state. Extracellular DNA is a ubiquitous structural component of biofilms that is stabilized by RuvA, and addition of the RuvABC complex results in significant biofilm disruption (Devaraj *et al*., 2019; Iwasaki *et al*., 1992). The oligopeptidase gene, *pepF*, was also downregulated in the biofilm state. PepF has previously been shown to influence biofilm formation in *Aeromonas hydrophila* (Du *et al*., 2016).

### Short-term adaptation and fixed genomic changes associated with biofilm formation involve distinct genes

We investigated the number of genes that were shared between the GWAS and differential gene expression analyses. Six genes exceeded the threshold for significance between the genome-wide association studies and differential expression analysis at 30°C and 37°C, respectively. No genes were identified as significantly associated with biofilm formation in all three experiments. Given the low numbers of shared genes observed between the experiments we sought to determine quantitatively whether this lack of overlap was statistically significant using a chi-square test. A chi-square test showed there is a significant relationship between the genes identified by the differential expression analysis and by GWAS at 30°C, χ^2^ (2, *N* = 2864) = 5.85, *p* < 0.025, and 37°C, χ^2^ (2, *N* = 2864) = 5.85, *p* < 0.001. The number of genes shared by the two types of experiment is less than would be expected through random sampling. This suggests that short-term adaptation to biofilm formation, investigated by differential gene expression, and long-term adaptation to biofilm formation, investigated by GWAS and involving fixed genomic changes, involve distinct genes (**Figure 5**).

**Figure 5:**
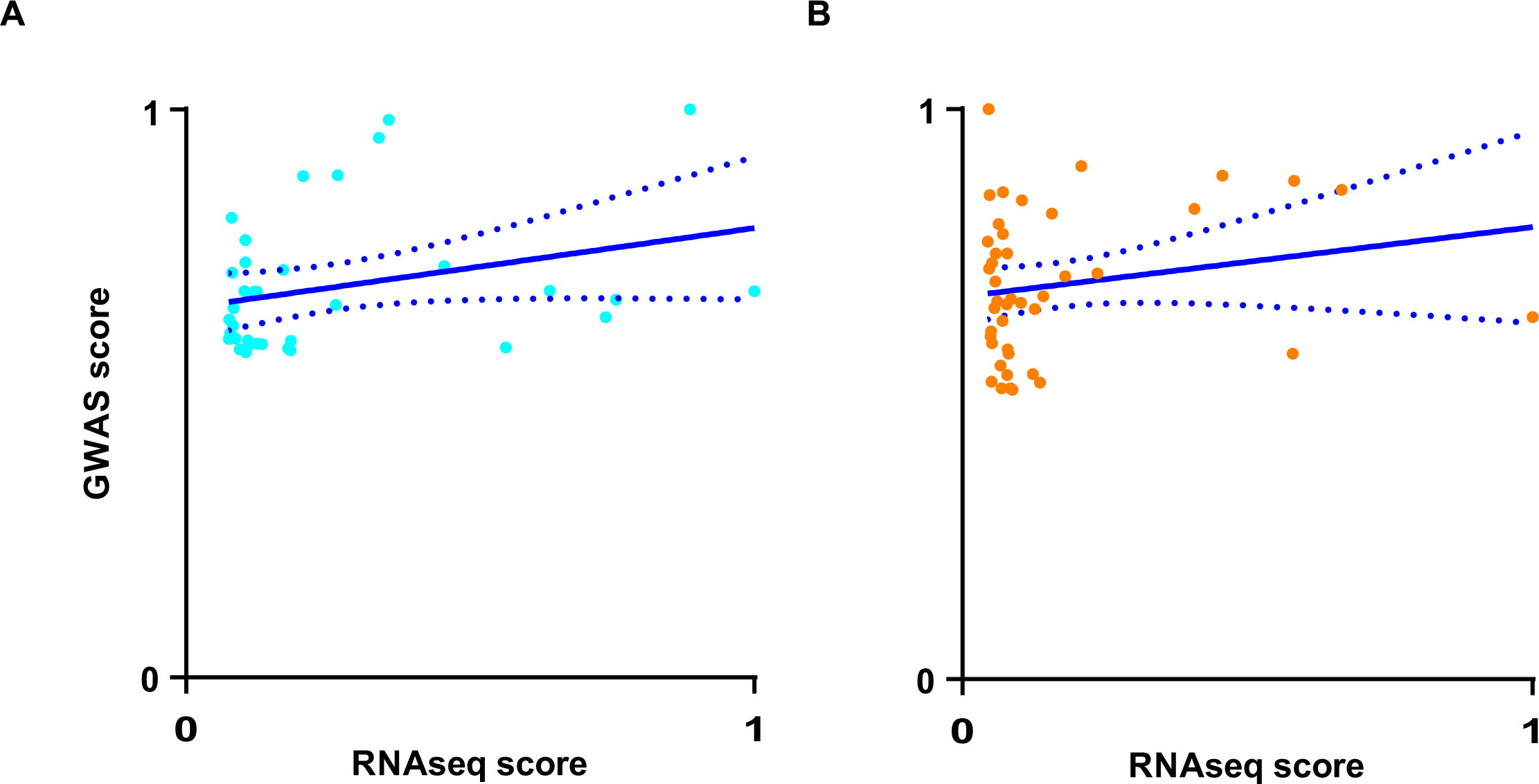
Biofilm adaptive changes in gene expression do not correlate with mutational changes identified by GWAS. Scatter plots of overlapping hits exceeding the suggestive significance threshold in genome-wide association studies and with a log(change) in gene expression greater than 2. GWAS scores were determined as the proportion of the maximal – log(*p*-value) observed per experiment. RNA-seq scores were determined as the proportion of the maximal log(change) observed. Colored dots indicate biofilm formation at 30°C (blue) and 37°C (red). The blue line indicates the line of best fit and dotted blue lines indicate 95% confidence intervals.

## Discussion

Comparing *L. monocytogenes* isolates from Japan with a large and geographically diverse collection revealed a diverse and highly structured population representing the major *L. monocytogenes* lineages circulating globally. The comparison allowed identification of genetically related isolate clusters that are internationally distributed suggesting recent, and possibly frequent, transmission between countries. Limited metadata associated with publically available isolate genome, made it difficult to draw conclusions on epidemiological links between related international clusters. However, these observations highlight the importance of real-time monitoring and tracking of *L. monocytogenes* genotypes at the international level, to identify common transmission sources.

The mixed ecological sources of samples from Japan allowed isolates to be classified according to their prevalence in different agricultural hosts, persistence in the food production environment or clinic. Putative differences in the isolate ecology can be inferred from the different clonal complexes sourced from each site. For example, isolates sourced from clinical samples are more likely to belong to the same complexes as isolates sourced from meat products. Alternatively, complexes isolated from agricultural samples are rarely sampled from other settings, except in the case of CC1, CC6 and CC7, which include isolates sampled from all three settings. However, the evolutionary events responsible for *L. monocytogenes* transmission between these settings remain poorly understood.

*In vitro* investigation of shortCterm diversification and selection for mutations associated with biofilm formation has revealed convergent mechanisms of evolution, with strains of the same species often presenting identical nucleotide changes (McElroy etCal., 2014). Furthermore, several genes are known to be associated with biofilm formation in multiple species of bacteria, for example, *luxS* has an important role in quorum sensing and is involved in biofilm formation in multiple species including *L. monocytogenes* (Sela et al., 2006), *Escherichia coli* (Niu *et al*., 2013), *Streptococcus mutans* (Merritt *et al*., 2003) and *Helicobacter pylori* (Cole et al., 2004). Importantly, in *L. monocytogenes* the ability to form a thick biofilm was distributed across the global phylogeny. This divergent origin of biofilm associated genes is consistent with multiple independent emergences of this important phenotype, potentially associated with HGT as observed in lineage II isolates from a relatively broad range of environments (den Bakker *et al*., 2008).

Selection through food production environments might be assumed to favour isolates that form thick biofilms and can survive on contaminated food to infect people. However, in this study, the mean biofilm forming ability of isolates was greatest amongst isolates sampled from agricultural sources compared with clinical sources or meat products. This observation likely reflects that heterogenous nature of biofilm formation in *L. monocytogenes,* limited sample numbers, and the importance of other factor in human infection.

We found a positive correlation between biofilm forming ability at the two temperatures investigated, with a significantly greater thickness observed at 30°C compared with 37°C. These findings are consistent with previous research on the influence of temperature on biofilm formation by *L. monocytogenes*, which suggests that biofilm formation varies with temperature, with optimal biofilm formation being observed at 30°C (Di Bonaventura *et al*., 2008)(Govaert *et al*., 2018). Non-motile mutants of *L. monocytogenes* are not capable of forming biofilms, indicating the importance of flagellar based motility in the formation of biofilms (Lemon, Higgins and Kolter, 2007). In fact, the reduction in biofilm formation between 30°C and 37°C could be a result of the temperature dependence of these structures involved in cell motility (Gründling *et al*., 2004). In this context, while our differential expression and GWAS analyses identify candidate biofilm genes, it will be important to confirm the link between sequence variation with function (Kobras *et al.,* 2021), particularly when temperature can influence biofilm formation (Di Bonaventura *et al*., 2008).

Adaptation can occur over different timescales. Long-term adaptation is typically considered in the context of alterations to genes, resulting from mutations or recombination, that confer stable inheritable fitness advantages (Sheppard, Guttman and Fitzgerald, 2018). Relatively rapid adaptation can also occur, potentially involving changes in regulation of gene expression rather that DNA sequence. This can include transcription factors (Lebreton and Cossart, 2017), epigenetic modifications (Casadesús and Low, 2006), and phase variation (Manuel *et al*., 2015a). These modifications provide a reversible and dynamic responses to environmental change, which can inherited over multiple generations (Bervoets and Charlier, 2019). Biofilm adaptation has been linked to phase variation of the DNA methyltransferase (*Mod*) in *Haemophilus influenzae* and alters the expression of genes controlled by the regulon via site-specific methylation (Brockman *et al*., 2018). Similarly, epigenetic regulatory modification can lead to distinct biofilm morphologies, despite isolates being genetically identical (Harris *et al.,* 2016)(Chia, Woese and Goldenfeld, 2008).

The transition between planktonic and biofilm states is major phenotype change (Toliopoulos and Giaouris, 2023; He and Ahn, 2011; Schembri, Kjaergaard and Klemm, 2003). Under the assumption that the most beneficial adaptations will increase most rapidly in the population, one might expect genes that exhibit changes in expression will also accumulate sequence variation linked to biofilm formation. However, in our study there was little evidence of overlap between biofilm genes linked to rapid and longer-term adaptation. This is consistent with a more complex adaptive process involving variation in both expression levels and DNA sequence. For example, observations of short– and long-term exposure to antibiotics have revealed initial adaptation through differential gene expression, including upregulation of efflux pumps (Händel *et al*., 2014). Subsequently, the SOS response promotes HGT and *de novo* mutation, resulting in resistance to higher antibiotic concentrations (Händel *et al*., 2014).

Understanding the contrasting processes linked to biofilm formation has particular significance for listeriosis. Broadly biofilms confer physical, adaptive or virulence advantages to the pathogen. First, physical biofilm barriers promote survival in food production environments and offer protection against antibiotics prescribed to treat infections (Serra and Hengge, 2014). Second, the accumulation of DNA – including mobile genetic elements, within biofilms facilitates HGT adaptation to changing environmental conditions. Finally, as in many pathogenic bacteria, biofilms are linked to virulence. There is clear utility in understanding contrasting genetics underpinning biofilm adaptation. Mature biofilms are linked to persistent infections and are more difficult to treat with antibiotics so early interventions are beneficial. Considering both gene expression and sequence polymorphism as important in biofilm formation may help in understanding important pathogenic lineages and guide interventions to reduce their emergence and spread.

## Author statements

### Contributors

SKS and BP designed the study, and wrote the paper with WM. WM, EM, JC, MH, BP, KY and HA performed experiments and analyses. MDH sequenced the isolates. NM helped with statistical analysis. All authors contributed and approved the final manuscript.

## Acknowledgements

We acknowledge Dr Susan Murray and Dr Rhys Tancock-Jones (both formerly Swansea University Medical School) for help extracting DNA for sequencing.

